# Neev, a novel long non-coding RNA, is expressed in chaetoblasts during regeneration of *Eisenia fetida*

**DOI:** 10.1101/806661

**Authors:** Surendra Singh Patel, Sanyami Zunjarrao, Beena Pillai

## Abstract

*Eisenia fetida*, the common vermicomposting earthworm, shows robust regeneration of posterior segments removed by amputation. During the period of regeneration, the newly formed tissue initially contains only undifferentiated cells but subsequently differentiates into a variety of cell types including muscle, nerve and vasculature. Transcriptomics analysis, reported previously, provided a number of candidate non-coding RNAs that were induced during regeneration. We found that one such long non-coding RNA (lncRNA) is expressed in the skin, only at the base of newly formed chaetae. The spatial organization and precise arrangement of the regenerating chaetae and the cells expressing the lncRNA on the ventral side clearly support a model wherein the regenerating tissue contains a zone of growth and cell division at the tip and a zone of differentiation at the site of amputation. The temporal expression pattern of the lncRNA, christened Neev, closely resembled the pattern of chitin synthase genes, implicated in chaetae formation. We found that the lncRNA harbours 49 sites for binding a set of four miRNAs while the Chitin Synthase 8 mRNA comprises 478 sites. The over-representation of shared miRNA sites suggests that lncRNA Neev may act as a miRNA sponge to transiently de-repress chitin synthase 8 during formation of new chaetae in the regenerating segments of *Eisenia fetida*.

**Summary statement:** The earthworm, *Eisenia fetida*, regenerates posterior segments following amputation. The transcriptome of the regenerating worm revealed a novel lncRNA, expressed only at the base of regenerating chaetae. We propose that this lncRNA is a miRNA sponge that modulates chitin synthesis.

## Introduction

Earthworms are a large diverse group of segmented worms that inhabit niches just under or deep within the soil. The tube within a tube body plan of the earthworm comprises a muscular outer wall enclosing a gut within. It also has a simple vascular system to circulate blood and a nervous system comprising a nerve ganglion at the anterior end and a long ventral nerve cord running the length of the body and ring nerves within each segment.

Earthworms vary widely in their ability to regenerate. *Eisenia fetida* (commonly known as red wriggler) regenerates nearly 2/3rd of its posterior end (Xiao et al., 2011). The earthworm presents an invertebrate model of epimorphosis, a type of regeneration involving the restoration of original anatomy and polarity followed by de-differentiation, proliferation and differentiation of cells (Bely, 2014; Gazave et al., 2013; Planques et al.). Since each segment consists of nerve, muscle, vasculature and additional specialized structures, it provides a model for studying regeneration coordinated across different tissue types. For instance, chaetae, specialized projections embedded in the skin used for gripping the soil are controlled by nerves in each segment to achieve a well-coordinated crawl.

We have previously characterized the genome and transcriptome of the regenerating earthworm (Bhambri et al., 2018). Injury and loss of posterior 2/3rd of *Eisenia fetida* was followed by apparent wound healing in 5-10 days. A stub of tissue largely consisting of a mass of undifferentiated tissue was formed by 15 days and differentiated segments were formed by 20 days post-amputation. The period between 10 to 20 days after the injury presents a time window during which cell proliferation, growth and differentiation happens simultaneously in a 4-5mm long tissue amenable to molecular and cellular visualization.

The transcriptome of the regenerating worm revealed signatures of rapid cell proliferation, reorganization of extracellular matrix and the differentiation of nerves. Besides these signatures, we also reported the dynamic expression of non-coding RNAs that potentially play roles in regulating the timely, controlled and spatially organized transcriptome (Bhambri et al., 2018). Here, we report the expression pattern of selected non-coding RNAs in the regenerating earthworm. Besides validating our previous report, we focus on an lncRNA, which showed a unique expression pattern at the base of the chaetae.

Chaetae are stiff chitinous structures (appendages) that originate deep within the muscular body wall, the outer ends of which are used to grip the surface and in locomotor activity (Hausen, 2005). The lncRNA, named Neev, is expressed only in the few cells at the base of chaetae in newly regenerated segments close to the site of injury. Its expression pattern closely resembles that of chitin synthase and chitinases involved in the formation of chaetae. Most importantly, the position of chaetae and the expression pattern of Neev support a model wherein the rapidly proliferating cells are restricted to the tip while the zone of differentiation is established close to the injury site (Gazave et al., 2013). Notably, Neev is not expressed at the base of the pre-existing chaetae. Clearly the lncRNA is spatially restricted to a few cells and expressed transiently during the formation of chaetae, but it is not required for its maintenance or function.

## Materials and Methods

### Experimental conditions

*Eisenia fetida* earthworms were originally procured from farmers engaged in vermicomposting and has been subsequently maintained in a plastic tray and fed with plant matter in the laboratory at around 22°C for several years. No specific permissions were required for procuring earthworms. They are not included in lists of endangered species and do not come under animals requiring ethical approval. Medium sized worms were collected before the experiment, rinsed in tap water to remove any soil sticking to the surface and amputated as described in our previous paper (Bhambri et al., 2018). The site of amputation was at about 2/3rd of the body length from the anterior end, thus retaining about 60 segments. After amputation, the worms were maintained in a separate container but under similar culture conditions. Regenerating tissue and about 2-3mm of the adjacent tissue from the pre-amputated worms were collected for in situ hybridization.

### RNA isolation and probe designing

The regenerated earthworm was rinsed thoroughly in running tap water followed by autoclaved milli-Q water, regenerated and adjacent control tissue from 20-30 earthworms was collected in 1ml Trizol kept on ice, respectively at 15, 20- and 30-days post amputation. A homogenous cell suspension was made by grinding these tissues using a homogenizer and 200ul chloroform was added and shaken vigorously for 15 seconds. After incubation for 10 minutes at room temperature for phase separation, the mix was centrifuged at 10,000g, 15 minute, 4°C. The upper aqueous layer was separated and an equal volume of Isopropanol was added, incubated for 5 minutes, centrifuged at 10,000g for 10 minutes. The resulting pellet was washed thrice with 70% ethanol at 10,000g for 5 minutes each. The air-dried pellet was dissolved in 50ul nuclease free water. RNA (1ug) was used to make cDNA using Transcriptor High Fidelity cDNA Synthesis Kit (Roche #5081955001) and primers (FP, ATA TGG TAC CGT CTG CTC CCA GGG TTA G; RP, ATA TGC GGC CGC CTT GTG TCG AGT GTA TTC AAT TGC) designed to amplify full-length transcript.

### PCR cloning and Sanger sequencing

Gel extracted PCR product was cloned using TOPO™ TA Cloning™ Kit (Invitrogen #450640) as manufacturer’s protocol. Sanger sequencing was performed to confirm the sequence of the clone. In vitro transcription to synthesize the probe was performed using DIG RNA Labeling Kit (SP6/T7 #11175025910) with SP6 or T7 polymerase after linearizing the plasmid by using restriction enzymes. The probes were purified by using NucAway™ Spin Columns (#AM10070).

### RT PCR

Total RNA from tissue samples was used for cDNA synthesis (as above) primed by oligodT primer. Gene specific primers listed in supplementary information were used in quantitative RT-PCR reactions containing SYBR green master mix (Takara #RR820). The Ct values were used to calculate Fold Change against spike-in control lncRNA (see results for details) using the method described by Michael W. Pfaffl (Pfaffl, 2001).

### In situ hybridization

Regenerated earthworms were collected at 10, 15, 20- and 30-days post amputation (dpa) and washed thoroughly in running tap water followed by autoclaved milli-Q water. Earthworms were fixed overnight at 4°C in 4% (w/v) paraformaldehyde (PFA) prepared in 1x PBS. After fixation, they were washed stringently in PBST (0.1% Tween 20 in PBS) with subsequent storage in 100% methanol at 4°C. Prior to hybridization, the stored earthworms were rehydrated with gradient of 90%, 75%, 50%, 25% and 0% (v/v) methanol in PBST for 45-60 min each. Earthworms were permeabilized by 20ug/ml Proteinase K for 45 minutes at 55°C. Fixed them again in 4% PFA for 20 minutes, blocked using hybridization buffer (50% formamide, 1.3x SSC, 5mM EDTA, 5% Dextran sulphate, 0.2% Tween 20, 100ug/ml heparin & 50ug/ml yeast t-RNA in DEPC treated water) for 60 minutes at 65°C. Hybridization was performed using sense and antisense probes prepared in hybridization buffer overnight at 65°C in water bath. Stringent washes were performed at 65°C with hybridization buffer thrice for 30 minutes each followed by washes in TBST (0.1% Tween 20 in TBS) for 15 minutes at room temperature. After incubation at room temperature for 4 hours in 1:2000 dilution of anti-digoxigenin antibody (Roche #11376623) prepared in TBST containing 10% FBS, TBST washes of 15 minute each at room temperature were performed thrice and subsequently the tissue was stained using NBT (working concentration 500ug/ml; Roche #11383213001) and BCIP (working concentration 562.5ug/ml, Roche #11383221001) in developing solution (0.1M NaCl, 0.1M Tris.HCl pH 9.5, 0.05M MgCl2, 0.1% Tween 20 in DEPC treated water). Images were captured by mounting earthworms in 2.5% methylcellulose at 3.2x and 5x magnification.

## Results

On the basis of transcriptomics analysis in the regenerating earthworm, we prioritized sixteen potentially non-coding RNAs for further validation, since they were about 1kb or larger and free of low complexity repeats. We designed qRT-PCR assays to detect four of the predicted lncRNA using the assembled transcript sequences (see methods; Supplementary data 1). Although GAPDH is widely used as a control in gene expression studies, it was not suitable for normalization in our experiments because it is strongly upregulated during regeneration. Instead, we used a spike-in normalization method, by adding an in vitro transcribed RNA fragment to the qRT-PCR reaction. The spike-in control lncRNA was originally cloned from the zebrafish genome that has no sequence homology with the earthworm genome (Sarangdhar et al., 2017). We also verified that the primers for this fragment produce no product when provided with the earthworm cDNA as a template. In close agreement with the RNAseq data, all four lncRNAs were strongly over-expressed in the regenerating tissue (Figure 1). As shown in the figure, the newly regenerated segments expressed the lncRNAs at 4 to 8 times the levels in the adjacent control segments.

**Figure 1:**
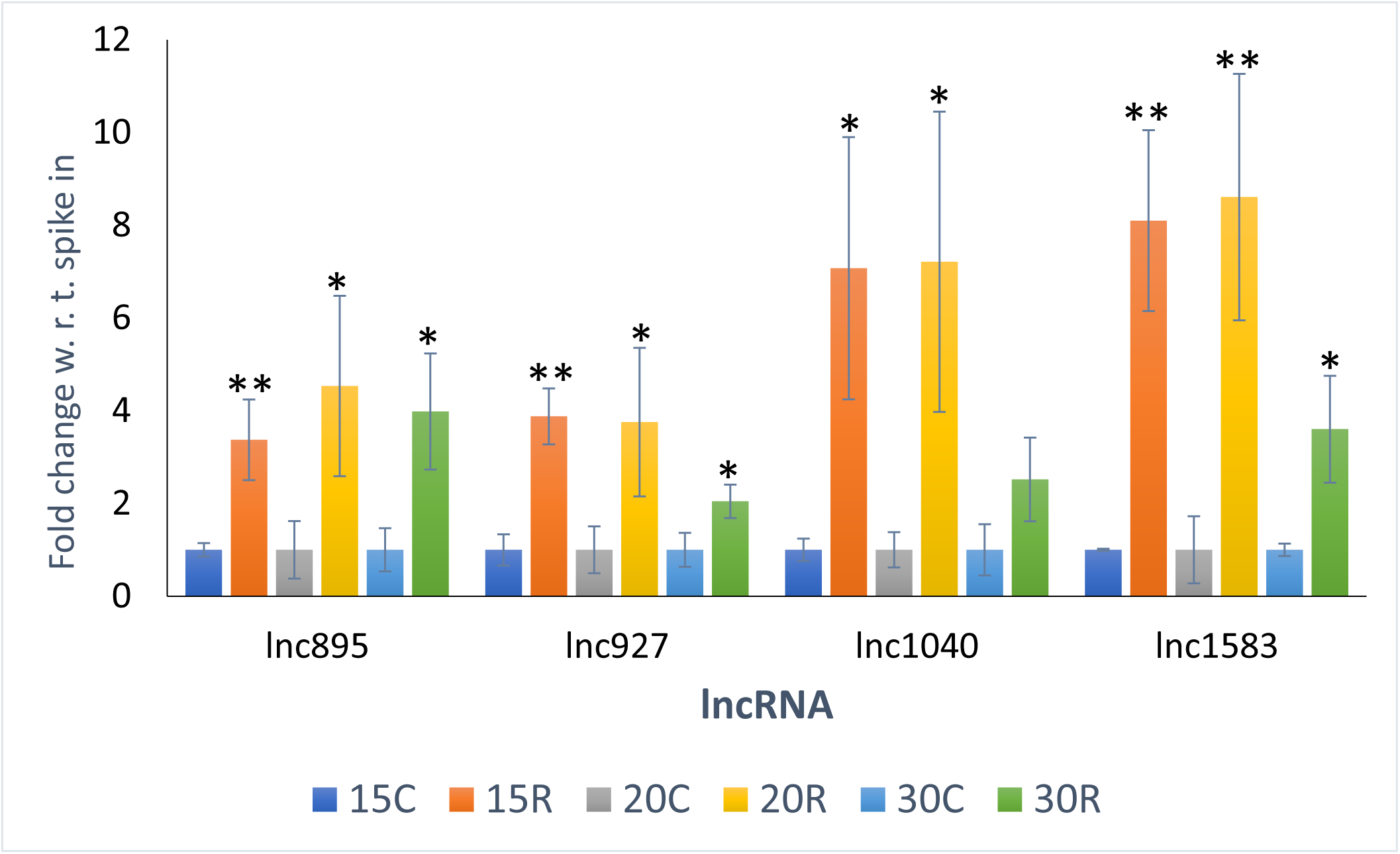
Validation of novel lncRNA upregulated during regeneration by qRT PCR. *Eisenia fetida* worms were amputated at approximately the 60th segment from the anterior end and allowed to regenerate over a period of 30 days. At 15, 20- and 30-days post amputation, the regenerating tissue (15R, 20R, 30R) and adjacent control tissue (15C, 20C, 30C) were collected. RNA isolated from these tissues were used for qRT-PCR (see methods for details; primer sequence). The lncRNAs, being novel were named according to the size of the transcript assembled from RNAseq data. (Biological replicates (N)=3; technical replicates (n)=3; t-test, * p-val<0.05; ** p-val<0.01)

Next, we cloned each lncRNA gene into the TOPO TA cloning vector and generated digoxigenin labelled probes by incorporating digoxigenin linked rUTP in the in vitro transcription reaction. These probes were used in in situ hybridizations in the collected sample containing regenerating tissue closely juxtaposed with the tissue from the worm before injury. Although there was a detectable signal in the regenerating tissue, the expression of the lncRNAs did not, in general, have a distinctive spatial pattern (Supplementary figure2). A notable exception was the lncRNA of 895nt length that showed a recurring pattern of expression with four spots in each segment on the ventro-lateral and ventral side in the regenerating region (Figure 2). Estimating from the Ct values, the lncRNA was expressed at about one-tenth the abundance of GAPDH, making it quite abundant, since lncRNAs are usually expressed at low levels. Since this abundant lncRNA was restricted to tiny spots, the expression was strong and clearly visible.

**Figure 2:**
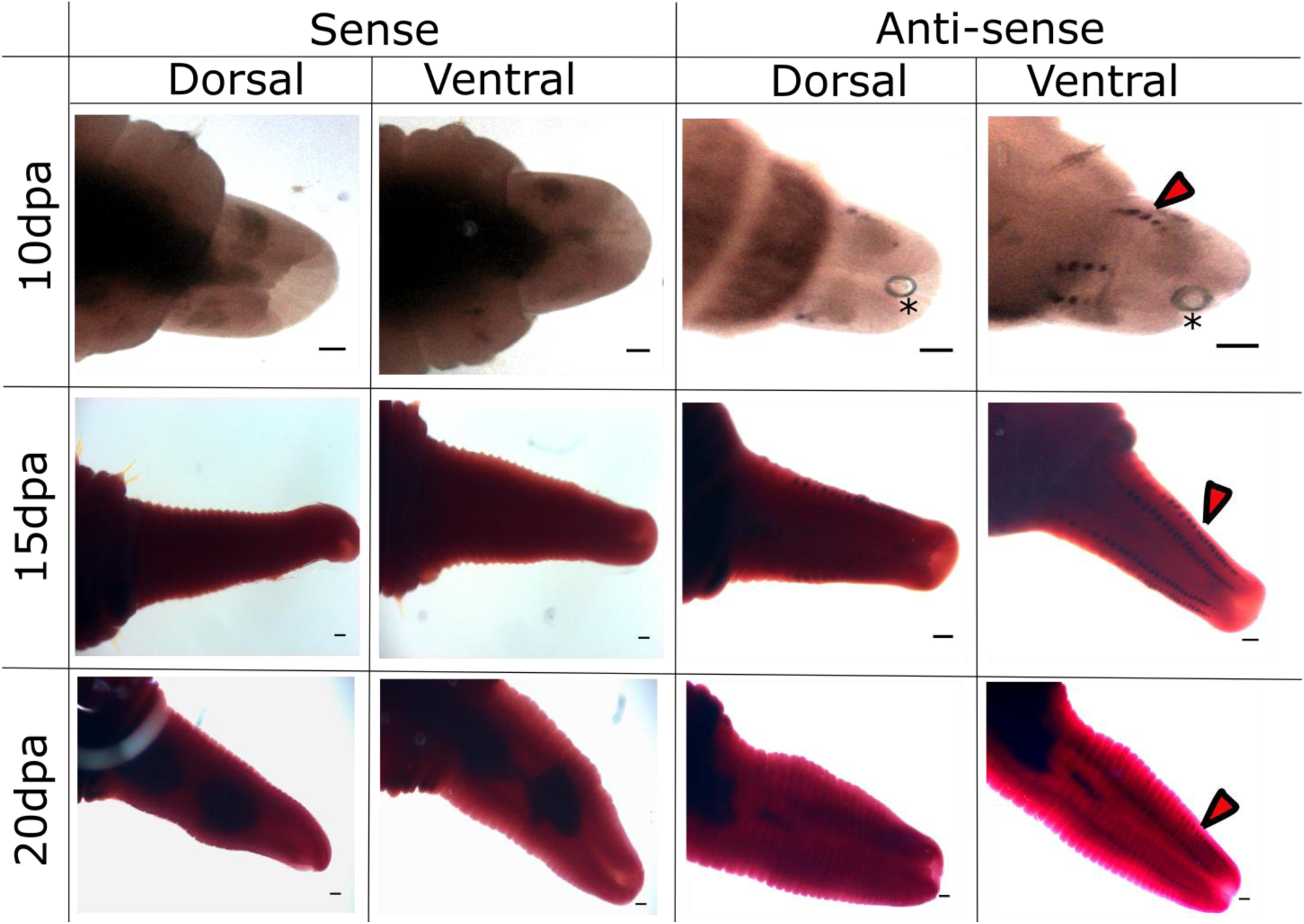
lncRNA Neev expression pattern in *Eisenia fetida* regenerating tissue. Earthworms were allowed to regenerate for 10, 15- and 20-days post amputation (dpa). Whole mount in situ hybridization of the regenerating tissue was performed using probe RNA synthesised by in vitro transcription. The antisense probe (right) showed distinct expression signal on the ventral side (arrowhead), while the sense probe (left) served as negative control (n=5; typical results are shown here). Asterisks marks are placed over air bubbles (Scale bar=100micrometre)

To understand the source of the signal (Figure 3A), we made longitudinal incisions on the dorsal region and spread out the inner body wall. By gently teasing out the tissue around the signal, it was clear that the spots were at the base of newly assembled chaetae (Figure 3B-C). Notably, no such spots were seen at the base of the chaetae before regeneration (Figure 3A). Clearly, this lncRNA was strongly but transiently induced in a very small group of cells closely associated with the chitinous setae. Due to this interesting expression pattern, we named the lncRNA Neev, which means base or foundation in Hindi.

**Figure 3:**
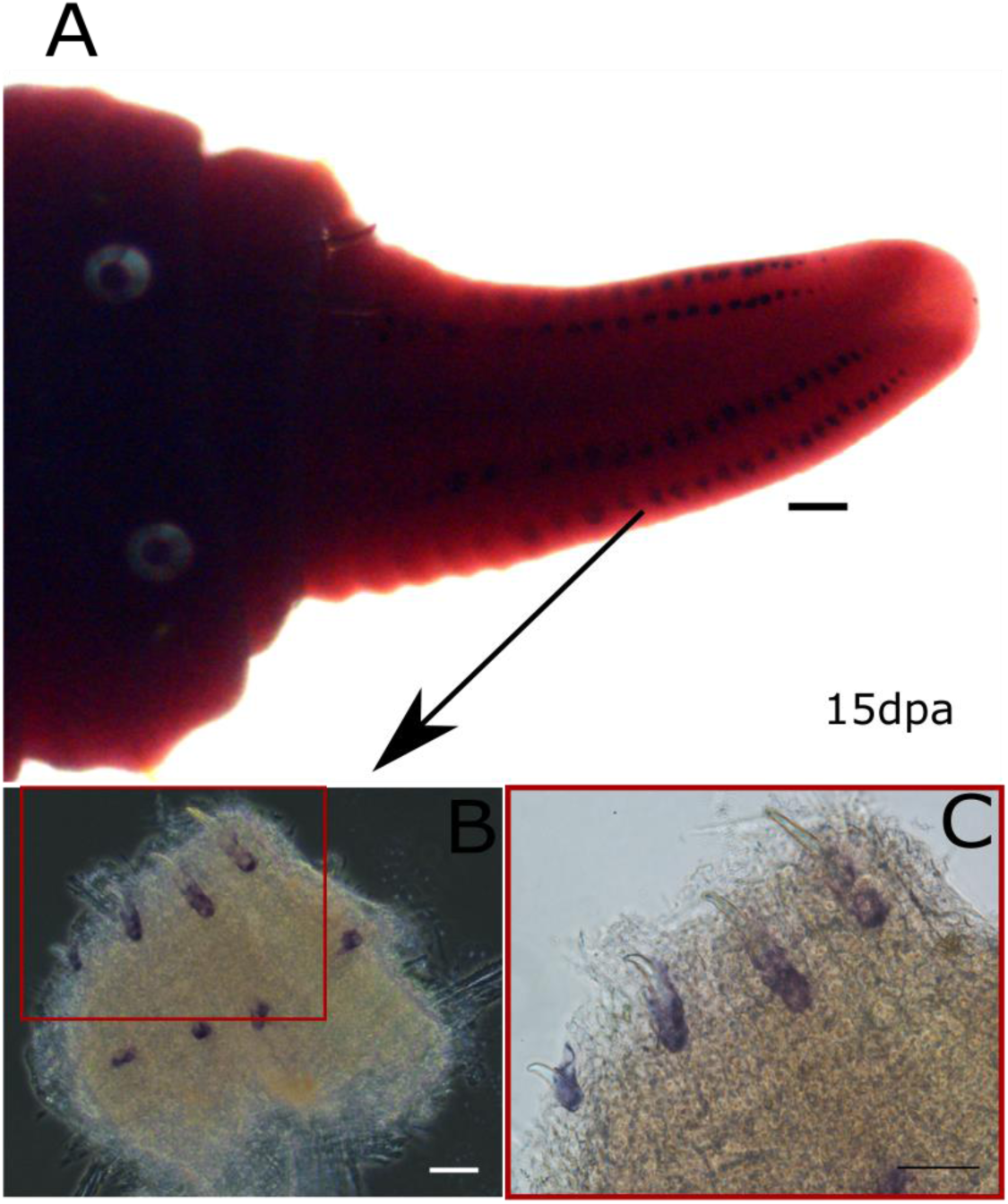
The lncRNA, Neev, is expressed at the root of chaetae in the regenerating tissue. After 15days post amputation (15dpa) whole mount in situ hybridization was performed on the regenerating worm (see Figure 2 and top panel; scale bar: 100micrometre). The tissue was dissected from the dorsal side to expose the root of the chaetae and imaged at 10X magnification (B) and 40X to visualize the expression of the lncRNA.

In transcriptomics experiments, fragments of protein-coding mRNAs are sometimes erroneously annotated as non-coding transcripts. Some legitimate lncRNAs may also produce functional peptides from microORFs. To rule out spurious annotation and detect conserved microORFs, we aligned the sequence of the Neev lncRNA to genome scaffolds assembled previously. Although there was a 85aa open reading frame, it did not show any similarity to known proteins (Figure 4). In the scaffold from which the Neev gene was derived, we found a neighbouring conserved region of 232nt with strong similarity to many genomes including vertebrate model systems like zebrafish (Figure 4; bottom right panel). By aligning the transcript sequence to the genomic sequence, we found that the Neev gene comprises two exons and a 191nt long intron (Figure 4; bottom left panel). Taken together, Neev codes for an lncRNA that satisfies all the currently prevalent criteria, i.e. multi-exon, polyadenylated transcript longer than 200nt and is devoid of ORFs larger than 300nt (Clamp et al., 2007; Dinger et al., 2008; Frith et al., 2006; Niazi and Valadkhan, 2012).

**Figure 4:**
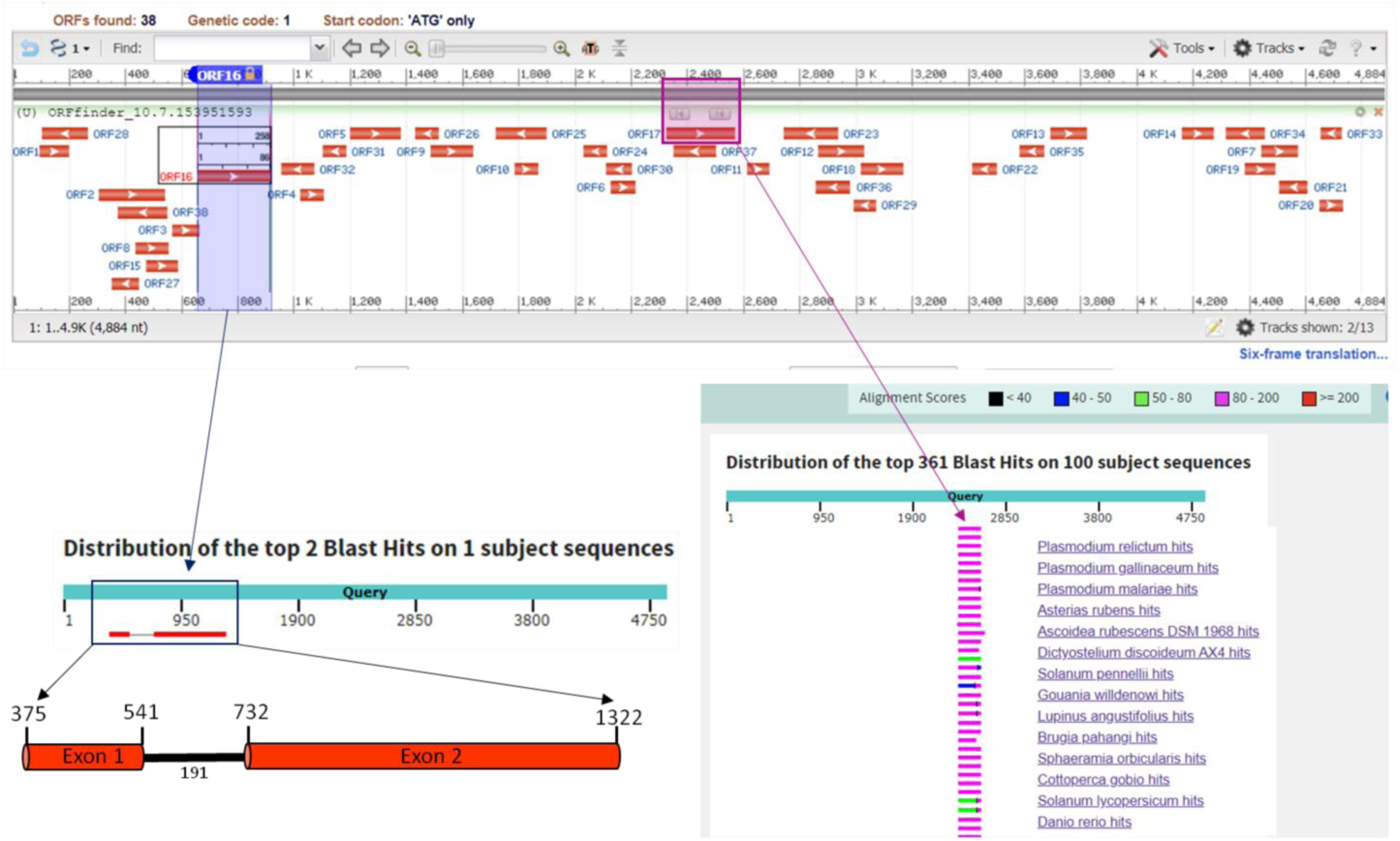
Alignment of lncRNA Neev with *Eisenia fetida* genomic scaffold SACV01159372.1. Prediction of Open Reading Frames (ORFs) in the genomic contig showed 38 small ORFs with no functional annotation (top panel). Neev lncRNA contains two exons (bottom left panel) and a 191nt intron. Only one ORF about 900nt in the 3’ direction was conserved in other organisms (bottom right panel).

Next, we tried to assign a potential function to the transcript. Since lncRNAs often regulate overlapping genes or genes in close proximity by RNA-DNA hybridization and recruitment of chromatin modifiers, we first checked for relevant ORFs in the 5kb contig containing the Neev gene. Since there were no genes in this region, we speculated that the lncRNA might regulate expression of distant genes through RNA-RNA binding. Chaetae, i.e. chitinous setae originate in bulbous cells called chaetoblasts, which put out microvilli that are subsequently coated with large amount of chitin, presumably produced within these cells (Schweigkofler et al., 1998). We reasoned that the chaetoblasts would also need to express chitin synthase genes transiently and in a highly regulated manner. We checked our transcriptomics data for the expression pattern of chitin synthase genes and chitinases. Amongst 29 genes with the word chitin in their name, 10 changed in expression during regeneration. We retrieved the basal expression level of these genes and in agreement with our prediction, one of the chitin synthases, Chitin Synthase 8 (Q4P9K9), showed a strong induction of 11 to 23-fold in the regenerating tissue, compared to the adjacent control tissue (Figure 5).

**Figure 5:**
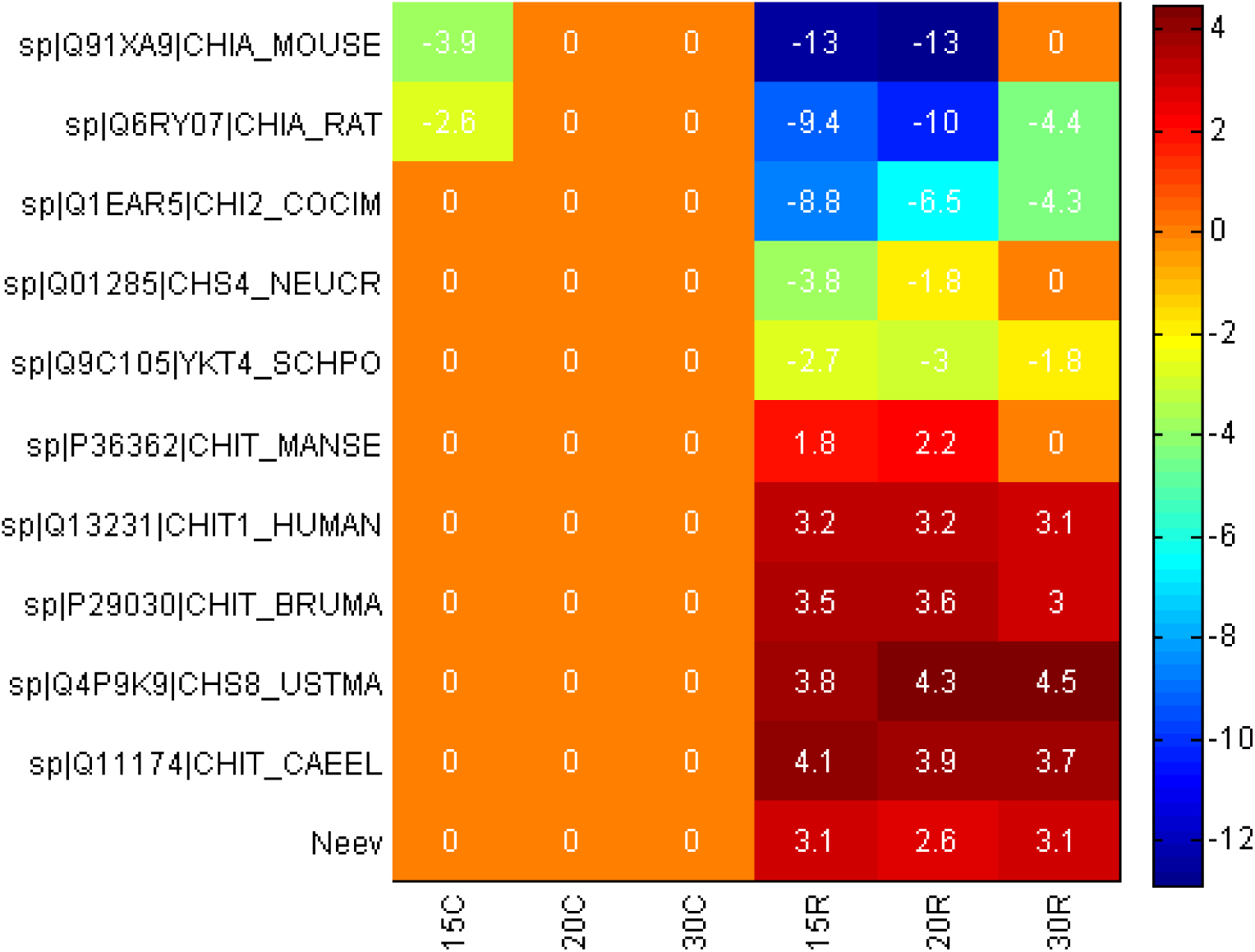
lncRNA Neev expression coincides with several chitin metabolism genes. The pre-amputated zone at 15, 20- and 30-days post amputation (15C, 20C and 30C) does not show any expression of these genes while four Chitin metabolism genes and Neev are strongly induced in the regenerating region at corresponding timepoints (15R, 20R and 30R). The numbers depict log2 Fold Change with respect to similar region of pre-amputated worm.

Next, we looked for potential RNA-RNA interactions that implicate the lncRNA in chitin synthase regulation. We aligned the sequences of lncRNA with differentially expressed chitin synthases. If they show similarity in anti-sense orientation, it could potentially form lncRNA-mRNA duplexes that are usually targeted for degradation. More complex regulatory mechanisms like guidance of splicing or RNA-RNA scaffold formation are also possible. In anti-sense orientation, the mRNA of Chitin Synthase 8 could potentially bind to the lncRNA only at four stretches of 7nt each.

We also looked for potential miRNA sponge like activity in the lncRNA sequence, because sequestration of miRNAs may transiently de-repress chitin synthesis genes to facilitate regeneration of chaetae. Since miRNAs tend to be highly conserved, we used the list of mouse miRNAs to predict targets and later verified that the earthworm genome contained sequences corresponding to the miRNAs of interest. We used the well-accepted miRNA target prediction tool, miRanda (Betel et al., 2010) to identify the most frequently occurring miRNA targets in Neev (Table 2). Two miRNAs were discarded from further analysis because we could not find the corresponding region in the earthworm genome. Four miRNAs, each with more than 5 target sites in the Neev lncRNA (see supplementary data2; Figure 4A) collectively had 49 sites of delG<-20C in the Neev lncRNA of 895nt length. Next, we checked the Chitin Synthase 8 mRNA sequence (length = 6202nt) to check for binding sites against the miRNAs. The four selected microRNAs had >100 sites on average and collectively had 478 potential binding sites (delG<-20C). To check for the rate of false positive predictions by the algorithm, we ran similar predictions for miRNA binding sites in three endo-chitinase genes, but none of them showed a comparable enrichment for binding sites of these miRNAs (Table 2). We also used 15 randomly selected 6kb regions (matching the length of Chs8 mRNA) from the earthworm genome, to ensure that the high number of targets was not a consequence of the length of the Chitin Synthase mRNA. By comparing the number of sites predicted in the randomly selected controls, we conclude that binding sites for the miR-667-5p and miR-7658-5p are over-represented in the Chitin Synthase 8 mRNA and the Neev lncRNA at a frequency significantly higher than expected by chance (pVal<0.001).

**Table 2:**

|  | No. of binding site |  |  |  |  |
| --- | --- | --- | --- | --- | --- |
| miRNA name | Neev | Chitin synthase<br>(DN356441 \| 6602nt*) | Endochitinase<br>(DN356441 \| 2488nt*) | Endochitinase<br>(DN319003 \| 1930 nt*) | Endochitinase<br>(DN331237 \| 2507 nt*) |
| mmu-miR-667-5p | 17 | 133 | 45 | 36 | 42 |
| mmu-miR-7658-5p | 16 | 130 | 35 | 28 | 43 |
| mmu-miR-7222-5p | 11 | 118 | 39 | 28 | 42 |
| mmu-miR-344-5p | 5 | 97 | 29 | 25 | 39 |

\*Trinity ID | length of mRNA (nt)

## Discussion

Long non-coding RNAs are a large group of transcripts that are more than 200 nucleotides in length, with no or some peptide coding potential with no well accepted criteria (Fang and Fullwood, 2016). LncRNAs are known to fold back into complex structures and mediate diverse functions in cells ranging from recruiting chromatin modifiers to guiding alternative splicing and sequestering microRNAs and proteins (Fernandes et al., 2019). We have previously identified several non-coding RNAs that were highly induced in the regenerating region of the earthworm from RNAseq data (Bhambri et al., 2018). The transcriptomics data was used in de novo assembly of transcript sequences, which were further classified as non-coding if they did not contain an open reading frame of more than 300 nucleotides in length. In agreement with the RNAseq study, the qRT-PCR validation also showed that the lncRNAs were induced in the regenerating region of the earthworm. The level of induction increased with time, till about 20 days post amputation but then tapered off. This coincides with the timeframe when the regenerated segments acquire almost all the features of the original segments and the transcriptional profile is largely restored to the normal pattern.

To the best of our knowledge, this lncRNA has no ability to encode a protein. While many dubious lncRNAs are now thought to be spurious products of abortive transcription (Consortium and The FANTOM Consortium, 2005; Ebisuya et al., 2008; Struhl, 2007) the presence of two exons in Neev suggests that it is more reliable (Derrien et al., 2012). It is polyadenylated, and only expressed from one strand i.e. it does not overlap with a protein coding gene. Neither could we find any ORF in close proximity within the 5kb contig containing the gene for Neev. Translating the 895nt transcript in all potential reading frames did not reveal any peptide with even minimal homology to a known gene. Unlike the typical lncRNA, Neev is expressed at high levels, only about 1/10th of the abundant GAPDH mRNA. Comparing the expression levels from the RNAseq data, it appears that Neev is more abundant than the chitin synthase 8 gene. Taken together, the reliable expression, poor conservation and presence of multiple exons agrees with it being a functional non-coding RNA.

Non-coding RNAs are found in large numbers in every genome, frequently outnumbering their protein coding counterparts (Derrien et al., 2012). The transient and localized expression of Neev is in agreement with the view that they may be involved in establishing the fine spatio-temporal regulation of genes in the regenerating tissue. LncRNAs are known to regulate gene expression at various levels: by recruiting chromatin modifiers (Campos and Reinberg, 2009; Kanhere et al., 2010), directing splicing (Bernard et al., 2010; Tripathi et al., 2010), modifying stability of mRNAs by masking motifs within the mRNAs (Kung et al., 2013; Matsui et al., 2008) or sequestering miRNAs (Faghihi et al., 2010; Franco-Zorrilla et al., 2007). Some of these mechanisms inherently involve RNA-protein complexes, which cannot be predicted on the basis of RNA sequence. However, some of the mechanisms can be predicted from the sequence of mRNA and lncRNA. The direct binding of lncRNA and mRNA can be deciphered from stretches of apparent complementarity while similarity at miRNA binding sites indicate the possibility of miRNA sponge mechanism. On the basis of high frequency of binding sites for miR-667-5p and miR-7658-5p in the mRNA of Chitin synthase 8 mRNA and Neev lncRNA, we speculate that the transient induction of the lncRNA in a few cells at the location of the chaetae, temporarily releases chitin synthase mRNA from repression, and facilitates the development of chaetae in the regenerating segments (Figure 6). Further experiments are needed to test the predicted miRNA sponge like activity of the lncRNA in vivo.

**Figure 6:**
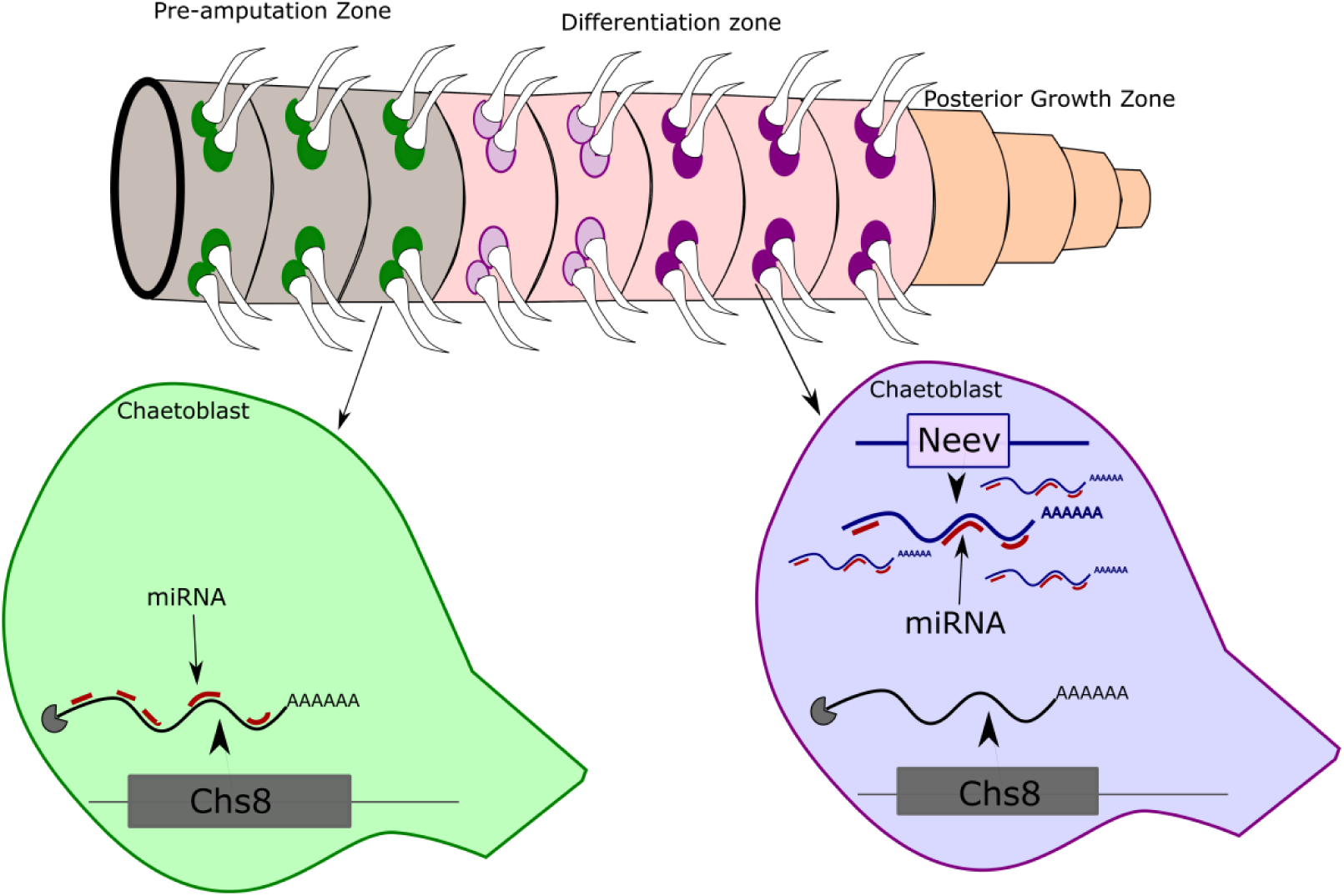
Schematic Depiction of gene regulation in the chaetoblast during regeneration. In the pre-amputation zone, Chitin Synthase 8 is repressed by miRNAs while in the differentiation zone, expression of Neev leads to sequestration of miRNAs and derepression of Chitin synthase 8. The posterior growth zone at the tip of the regenerating earthworm comprises undifferentiated cells.

The position of the regenerating chaetae and the pattern of Neev expression can help in inferring an invisible developmental boundary within the regenerating tissue. Immediately following the injury, a mass of undifferentiated tissue, called blastema caps the site of injury. Within days, the mass of cells starts differentiating to structures found within each segment even as rapidly dividing cells increase the volume of tissue. It was not clear if the temporal period of growth and cell division is completed before differentiation ensues. It is clear from our data that both differentiation and cell proliferation happen simultaneously. Further, it is also clear that the rapidly dividing cells are concentrated at the posterior tip while the differentiation zone abuts the site of injury. This is broadly in agreement with a recent report of zones within the posterior regenerating region of another annelid, Platynereis dumerilii (Gazave et al., 2013). Cells that exhaust their division potential are pushed down into the zone of differentiation where presumably local signals precisely position the chaetoblasts, which then adopt a transcriptional program distinctive from the neighbouring cells. At the site of injury, the newly formed segment has chaetoblasts expressing the Neev lncRNA, even as the pre-existing chaetoblasts (in the adjacent segment on the other side of the site of injury) remains immune to these local signals. The precise arrangement of the chaetoblasts in rows within each segment and columns that run across segments implies a very precise and local regulatory module that determines the expression of the Neev lncRNA.

## Acknowledgements

The authors acknowledge the support in genome analysis by Aksheev Bhambri. Technical support by Jayashree Niharika and Pooja Bharali who repeated the in-situ hybridizations reported here is gratefully acknowledged. Although the figures generated by them were not used, their independent validation of the data was useful in establishing the reliability of the data.

## Supplementary information

### Primers used for qRT PCR

Neev (lnc895) FP: GGTTCCAGAGCCGTAATGTTC

Neev (lnc895) RP: GCTACCATCATCGTCTTGCTG

lnc927 FP: TGAATGCTCGCTCTCCTACAT

lnc927 RP: TGGAATATTGCTGGAAAATGG

lnc1040 FP: ACCCAAAATGCACTGAACCAA

lnc1040 RP: TTTGCTCCGTCTTTTCGTCTG

lnc1583 FP: GAGCCAACTTGAACCAACTGT

lnc1583 RP: AATGCAACACTACCGACAACC

Spike-in qRT FP: CCTCTTGTATCTCACAGCTCAA

Spike-in qRT RP: CTGGAGACAATAGAAGGCAAT

### Supplementary data 1

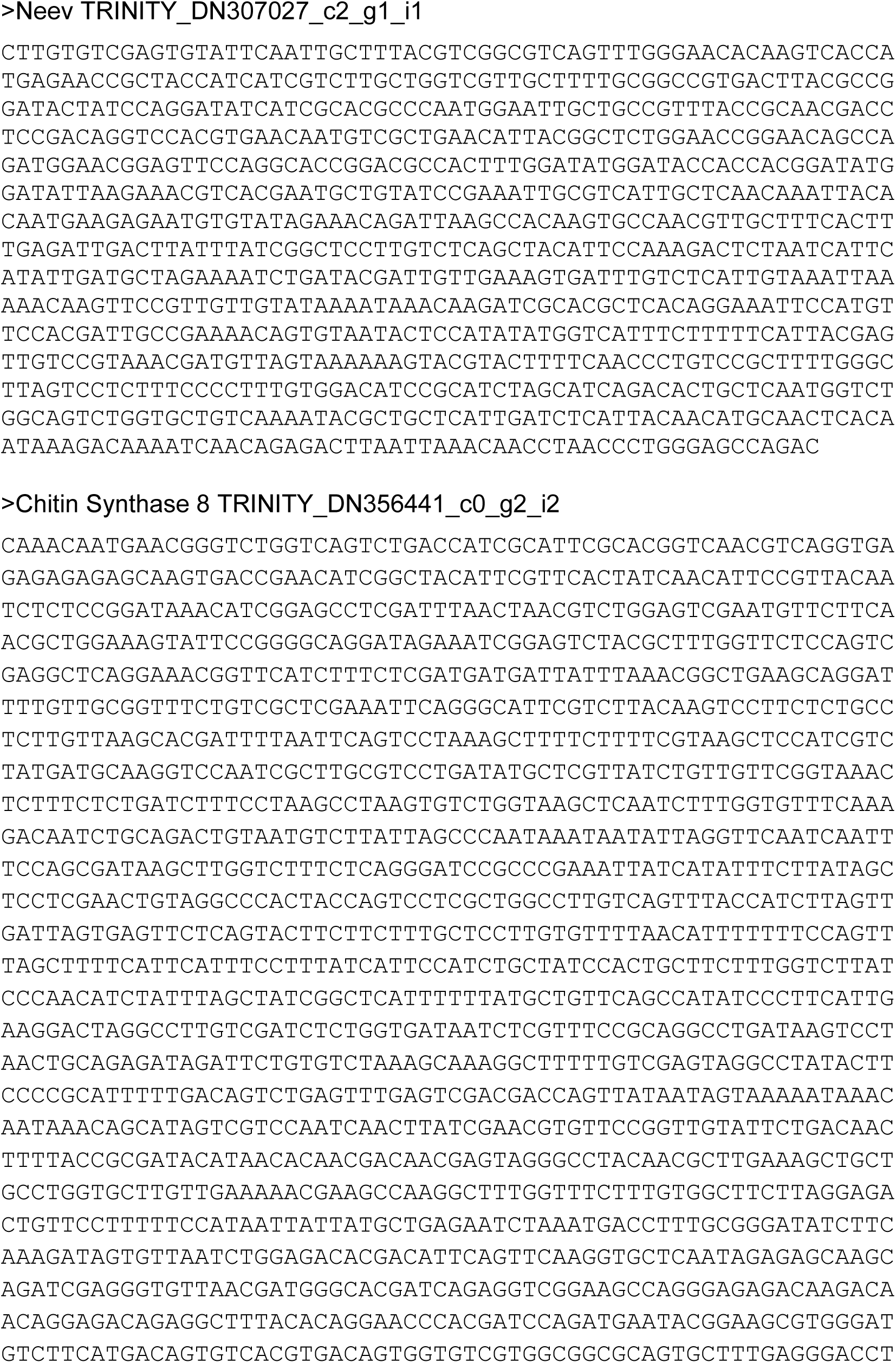
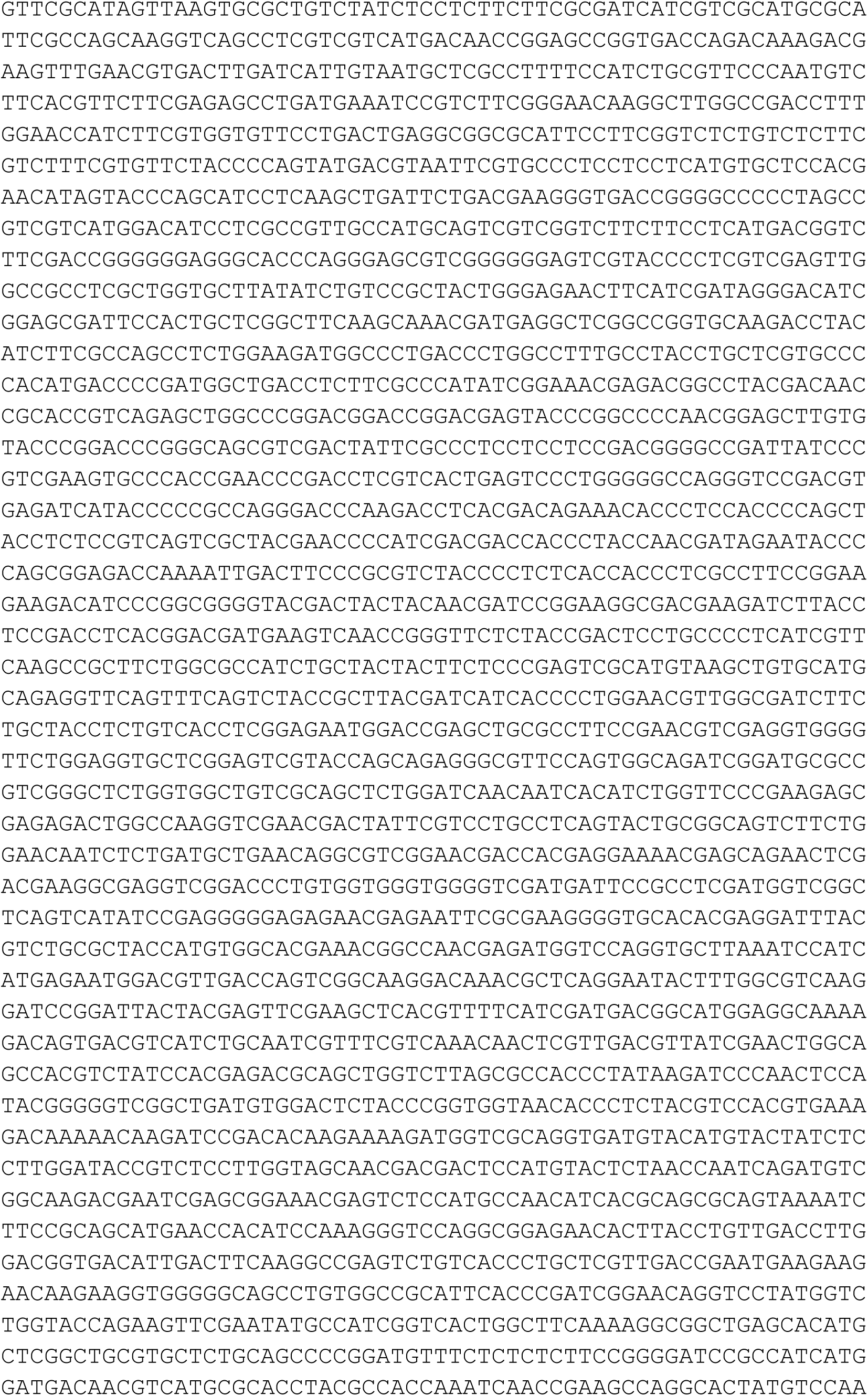
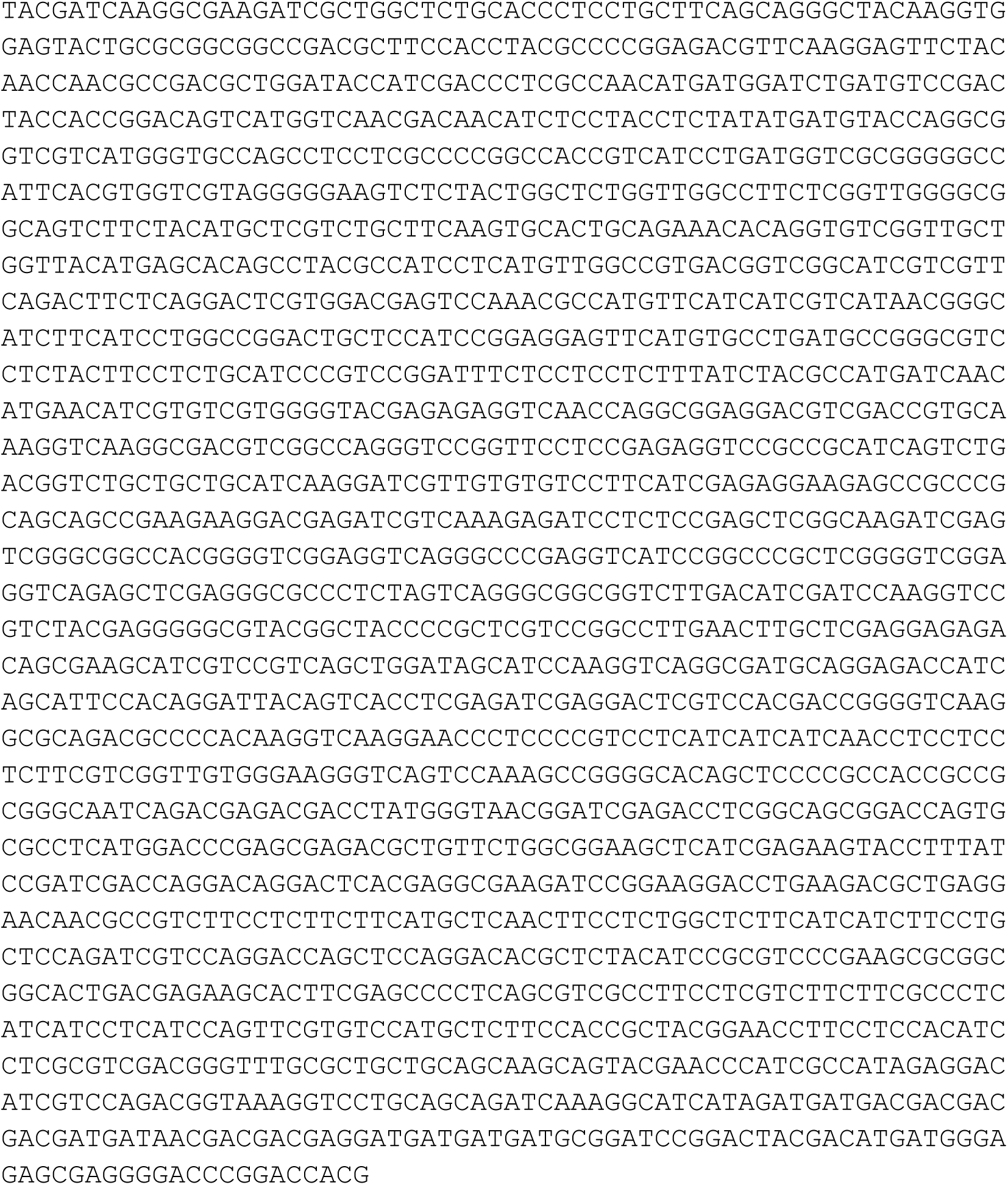

#### Supplementary data 2: Expression pattern of lnc927 in regenerating tissue

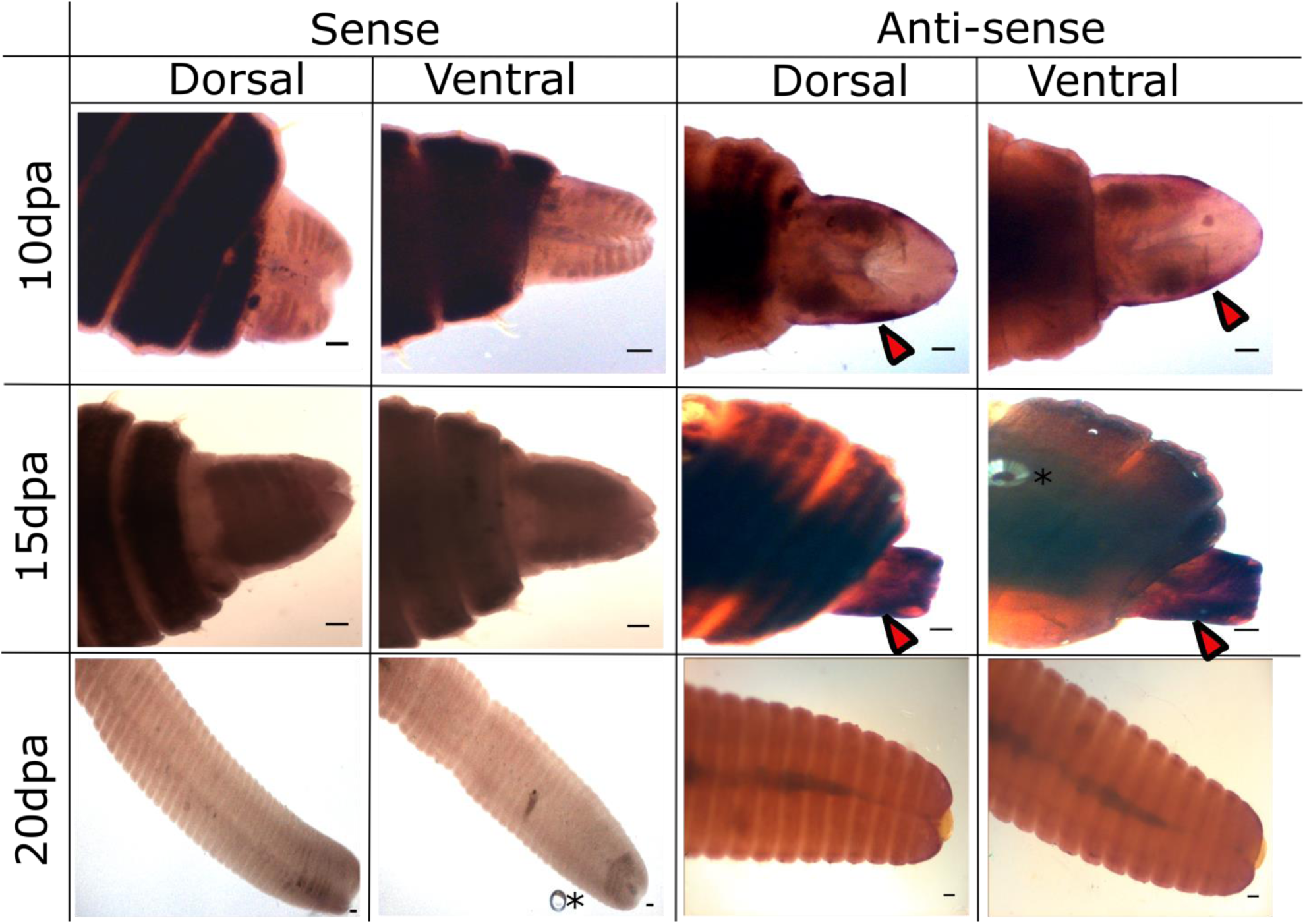

#### Expression pattern of lnc1040 in regenerating tissue

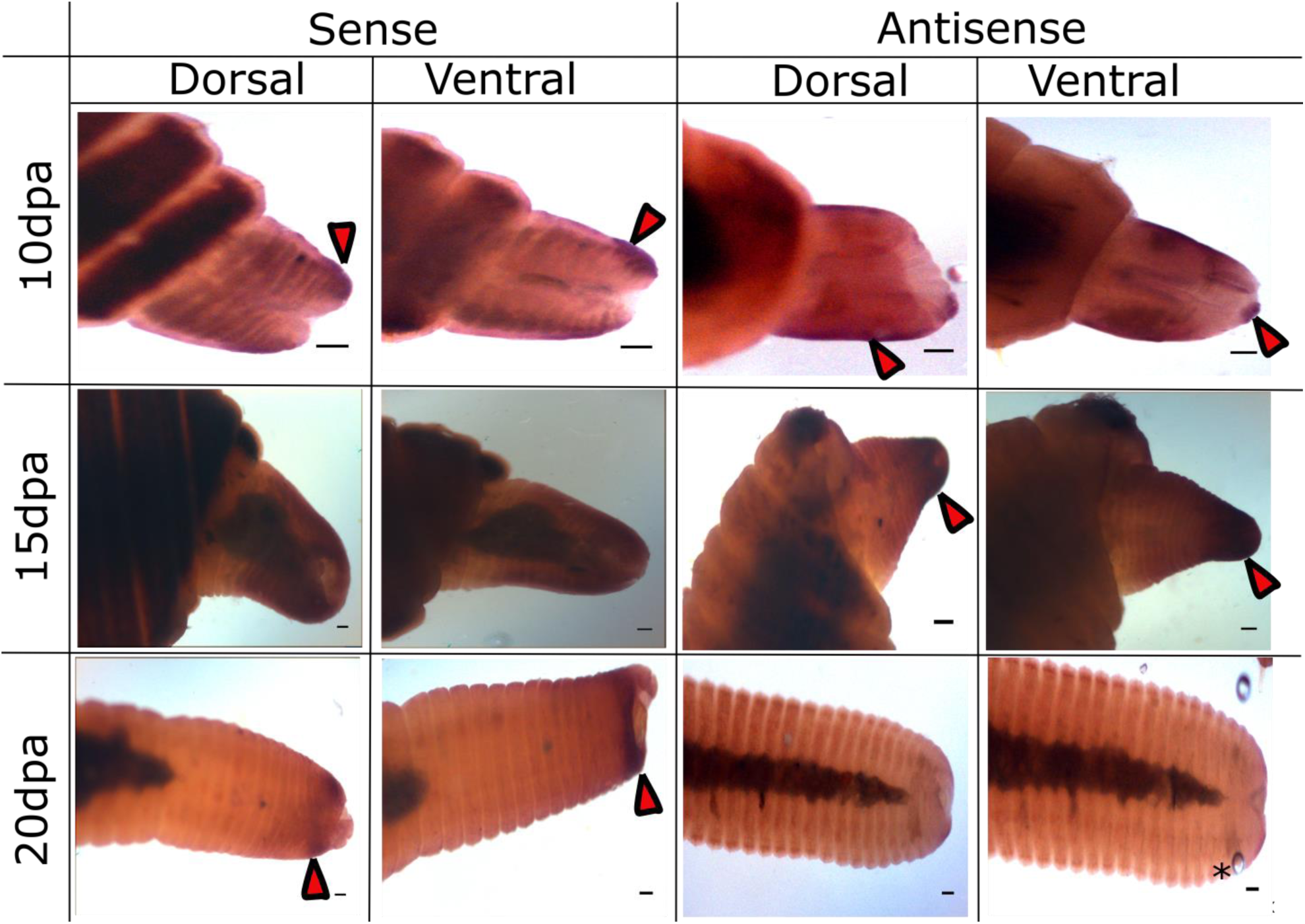

